# Across-speaker articulatory reconstruction from sensorimotor cortex for generalizable brain-computer interfaces

**DOI:** 10.1101/2025.10.08.680888

**Authors:** Ruoling Wu, Julia Berezutskaya, Zachary V. Freudenburg, Nick F. Ramsey

## Abstract

**Objective:** Speech brain-computer interfaces (BCIs) can restore speech features like articulatory movements from brain activity. However, for individuals with vocal tract paralysis, lack of articulatory movements can pose a challenge for speech BCI development. To address this challenge, our study aims at extracting generalizable articulatory features from a group of native Dutch speakers and reconstructing these features from brain data of a separate group of able-bodied individuals.

**Approach:** We applied a tensor component analysis (TCA) model to extract generalisable articulatory features from a publicly available articulatory movement dataset. To reconstruct articulatory features from the brain, we analyzed data of three able-bodied participants P1, P2 and P3 with high-density electrocorticography (HD-ECoG) electrode arrays implanted over the sensorimotor cortex. For each participant, a separate TCA model was applied to extract neural features. A gradient boosting regression model was used to reconstruct articulatory features from neural features. Reconstruction performance was measured as Pearson’s correlation coefficient (PCC) between the reconstructed and the generalizable articulatory features.

**Results:** The extracted articulatory features showed even contributions across participants, indicating that these features captured generalizable articulatory kinematic patterns. By using these features, we were able to reconstruct articulatory features from brain data. PCC between the reconstructed and original articulatory features were significantly above chance for all three participants, with mean PCCs of 0.80, 0.75 and 0.76 for P1, P2 and P3 respectively.

**Significance:** With the rapid development of speech BCI, our research demonstrates that speech-related articulatory features can be restored from HD-ECoG signal using generalizable articulatory features derived from able-bodied individuals. With the potential to reconstruct audio or speech-related facial movements from the reconstructed articulatory features, our framework may provide a new way for developing speech BCIs for people unable to produce mouth movements.

## Introduction

To produce speech, our brain coordinates the movement of vocal tract muscles, referred to as speech articulators: lips, jaw, tongue, and larynx (Browman & Goldstein, 1992; Chartier *et al*., 2018; Silva *et al*., 2024). Due to motor impairments caused by medical conditions such as amyotrophic lateral sclerosis (ALS) or brainstem stroke, some people may lose the ability to control their vocal tract muscles to produce speech. The loss of speech can lead to barriers in social life and substantially impact life quality (Walshe & Miller, 2011).

Brain-computer interfaces (BCIs) may be able to help these people by restoring their communication ability. This can be done, for example, by converting neural signals from brain areas involved in speech production to text on computer display or synthesized speech as has been shown in several individuals (Silva *et al*., 2024). However, it is unclear how generalizable these BCIs are across people. Also, these BCIs require a lot of data to train the decoding model. Most speech BCIs use signal from the ventral sensorimotor cortex (vSMC), where articulatory kinematics information is strongly encoded (Chartier *et al*., 2018; Mugler *et al*., 2018). Therefore, reconstructing the intermediate articulatory movement features is a logical approach for speech BCIs. Recent BCI applications have demonstrated successful speech synthesis from articulatory features reconstructed from limited neural signals (Anumanchipalli *et al*., 2019; Metzger *et al*., 2023). Accurate reconstruction of vocal tract and facial movement has also led to a successful implementation of a digital facial avatar that a person with motor impairment could use for enhancing communication and self-expression (Metzger *et al*., 2023). So far, research has been focusing on connecting brain signals to articulation in the same individuals who retain their ability to control their vocal tract muscles to some extent. However, people who suffer from severe vocal tract paralysis may completely lose the ability to control vocal tract muscles and cannot produce any speech-movement or sound (Duffy, 2012). For these people, no articulatory or acoustic data are available to train BCI decoding models.

For individuals unable to produce any speech movements, transfer learning could provide a solution for developing articulation-based speech-BCIs. Transfer learning applies knowledge learned from one dataset to a different but related dataset (Jayaram *et al*., 2016; Weiss *et al*., 2016). In the context of speech-BCIs, transfer learning can be used to extract articulatory features shared across healthy individuals and then map them onto the brain activity of individuals with vocal tract paralysis.

Two key research questions underlie the integration of speech-BCIs with transfer learning: (1) given individual variation in vocal tract anatomy, is it possible to extract generalizable articulatory features across healthy participants? (2) is it possible to reconstruct generalizable articulatory features from brain activity of a different group of individuals?

To address these questions, we developed a novel BCI framework that extracts generalizable articulatory features across healthy participants and reconstructs them from brain data collected from a separate group of participants without using their own articulatory movement data. We used a tensor component analysis (TCA) model to extract generalizable articulatory features (Kolda & Bader, 2009; Williams *et al*., 2018). We then used a gradient boosting regression model to reconstruct the articulatory features from high-density ECoG (HD-ECoG) data of three different participants, unseen by the TCA model. We show that the articulatory features can be reconstructed from HD-ECoG with a significant Pearson’s correlation with the original articulatory features, which demonstrates the possibility of cross-participants articulatory-brain mappings. Our result highlights the potential of the novel framework for developing generalizable BCI models that could facilitate speech reconstruction in people with vocal tract paralysis.

## Materials and Methods

### Electromagnetic articulography (EMA) dataset

We used articulatory movements recorded during word production from a publicly available Electromagnetic Articulography (EMA) dataset (Wieling *et al*., 2016). The EMA data was collected by a portable 16-channel device (WAVE, Northern Digital InC.) at a sampling rate of 100 Hz during a word production experiment. A total of 21 speakers of the Low-Saxon Dutch dialect and 19 speakers of the Central Dutch dialect participated in the word production experiment. During the experiment, participants performed two tasks. Task I was a picture naming task, in which participants were presented with 70 pictures of objects sequentially and were instructed to name the objects out loud in their dialect. Task II was a consonant-vowel-consonant (CVC) sequences reading task, in which participants were presented with 27 CVC sequences on screen sequentially and were instructed to read them out loud in the standard Dutch dialect (Figure 1a). Therefore, the stimuli included 97 Dutch words in total (70 object names and 27 CVC sequences). Stimuli in both tasks were repeated twice in random order. For both tasks seven EMA sensors were attached to jaw, lower lip (LL), right corner of the lip (right lip, RL), upper lip (UL), tongue tip (TT), tongue body (TB) and tongue dorsal (TD) positions, to track movements of the corresponding articulator along three dimensions: up-down, anterior-posterior and left-right directions (Figure 2). Positions in the left-right direction were excluded from all analyses because speech movement are typically symmetrical and provide little extra information. We confirmed this with an analysis of variance along each direction, where scores for the left-right direction for tongue was very low (Table S1). The timestamps with pronunciation onset and offset of each word were included in the dataset.

**Figure 1.**
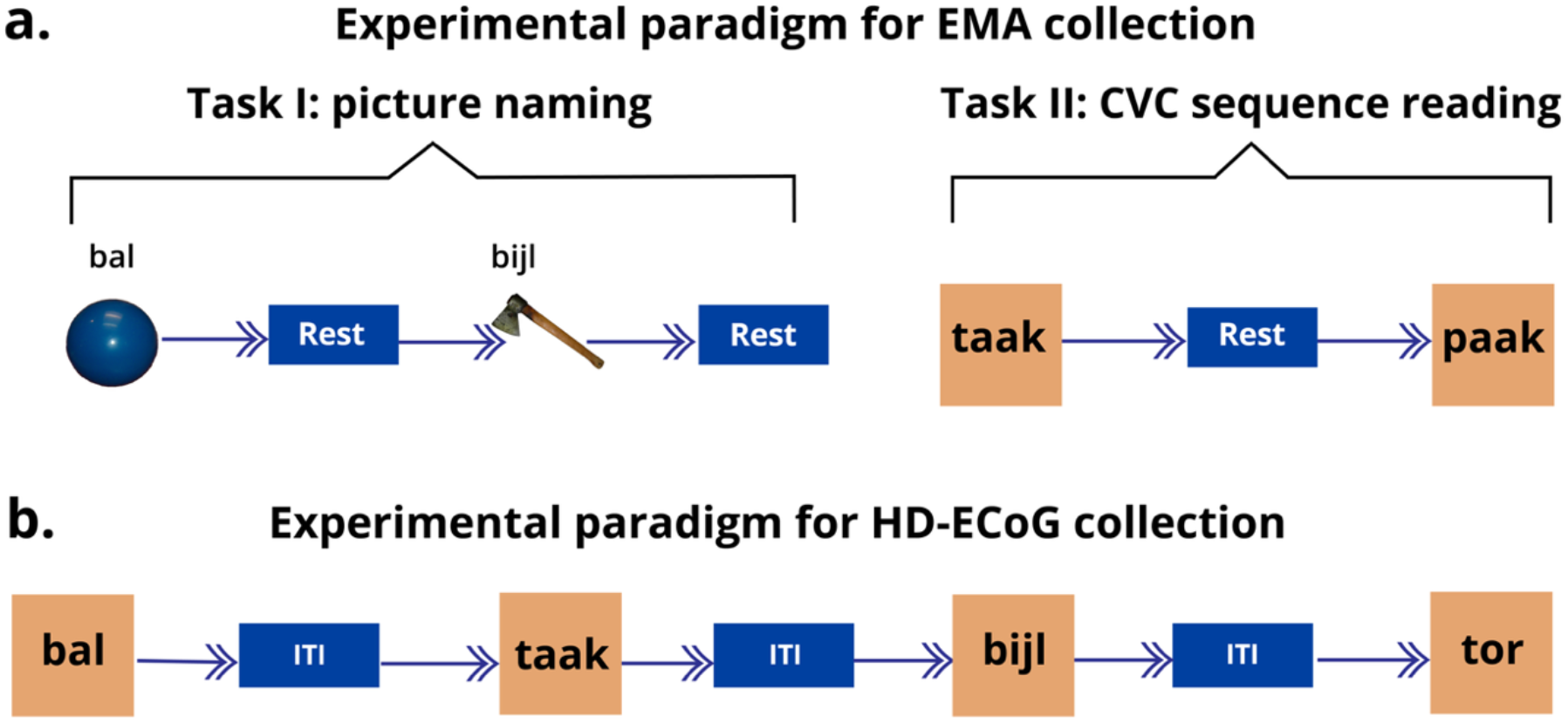
Experimental paradigms of EMA and HD-ECoG data collection. a. During task I of the EMA data collection, participants were presented with pictures of objects (70 in total) and were instructed to name them. During task II, participants were asked to read consonant-vowel-consonant (CVC) sequences (27 in total) presented on screen. Every picture/ CVC sequence was repeated twice in random order. This figure was adapted from images provided by Prof. Dr. Wieling, used with permission from the author. b. During the HD-ECoG data collection, participants were presented with words from the same set of 70 object names and 27 CVC sequences (97 Dutch words in total) used in the EMA experiment. Unlike the EMA experiment, all words were presented in textual form. The task trial duration was 2 seconds for P1 and P2, and 1.8 seconds for P3. The duration of inter-trial-interval was 1.2 seconds for P1 and P2, and 1.8 seconds for P3. Dutch words shown in the figure: *bal* (ball), *bijl* (axe), *taak* (task), *paak* (CVC letter sequence, not a real Dutch word), *tor*(beetle).

**Figure 2.**
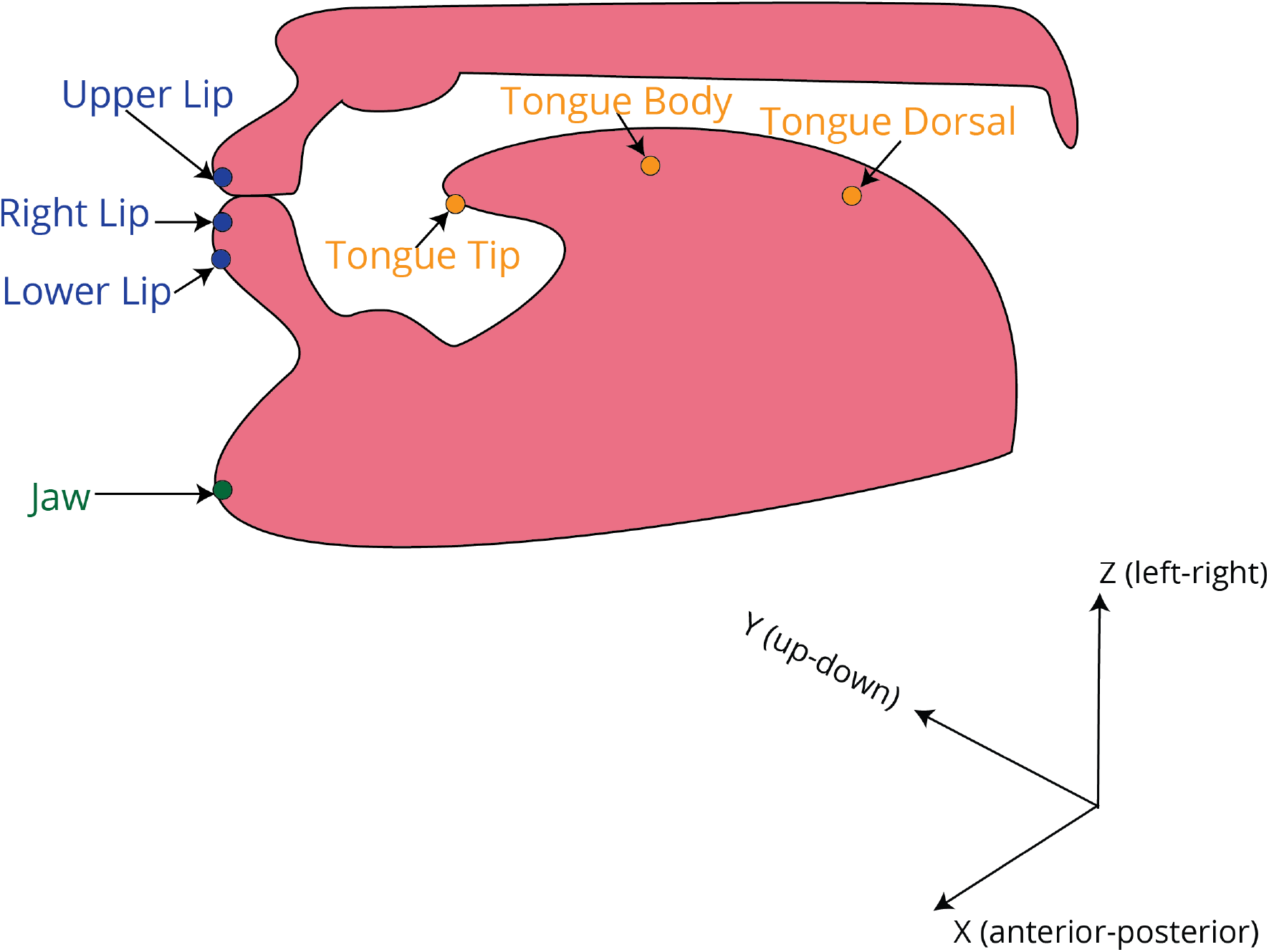
Sagittal view of locations of EMA sensors. Each sensor location is marked with a sphere, and the corresponding label is shown. There was one EMA sensor placed per location: jaw, lower lip, right lip, upper lip, tongue tip, tongue body and tongue dorsal locations to track movements of each corresponding articulator in three directions anterior-posterior (X-axis), up-down (Y-axis), and left-right (Z-axis) directions.

In our study, we only included data collected from participants speaking the Central Dutch dialect, given its greater similarity with the standard Dutch compared to the Low-Saxon Dutch dialect, as evidenced by a shorter Levenshtein distance (Heeringa, 2004). To ensure a consistent number of trials and comparable articulator movement amplitude, we additionally excluded data from two Central Dutch participants with missing trials, eight participants with interrupted trials and one participant with exceptionally large mouth movements (large amplitude of EMA signals beyond the range of mean±2.5*std). EMA data from eight remaining participants (U1 to U8) were included in our analyses.

### HD-ECoG dataset

#### Participants

Three male participants (P1, P2, P3, age 31, 26, and 40, respectively) were admitted to the University Medical Centre Utrecht for diagnostics and treatment of medication-resistant epilepsy or brain tumour. Two participants (P1 and P2) underwent clinical ECoG implantation for epilepsy diagnostic procedures and one participant (P3) underwent electrode implantation during awake tumour resection surgery. All participants provided written informed consent for implantation of a high-density ECoG grid for research purposes. The protocols were approved by the Medical Ethical Committee of the University Medical Centre Utrecht in accordance with the Declaration of Helsinki (2024). All participants consented to use of an high-density (HD) ECoG grid over the somatosensory cortex, which for P1 and P2 was implanted in addition to standard clinical grids with low density configuration. Here, we only focused on the HD-ECoG grids.

#### Data acquisition

All three participants were implanted with a HD-ECoG grid over the sensorimotor cortex on the left hemisphere (Figure 3), with different configurations (Table 1).

**Table 1.** HD-ECoG configurations across participants.

| Participant | Number of electrodes | Inter-electrode distance (IED) | Exposed diameter |
| --- | --- | --- | --- |
| P1 | 128 | 3 mm | 1 mm |
| P2 | 32 | 4 mm | 1 mm |
| P3 | 96 | 3 mm | 1 mm |

**Figure 3.**
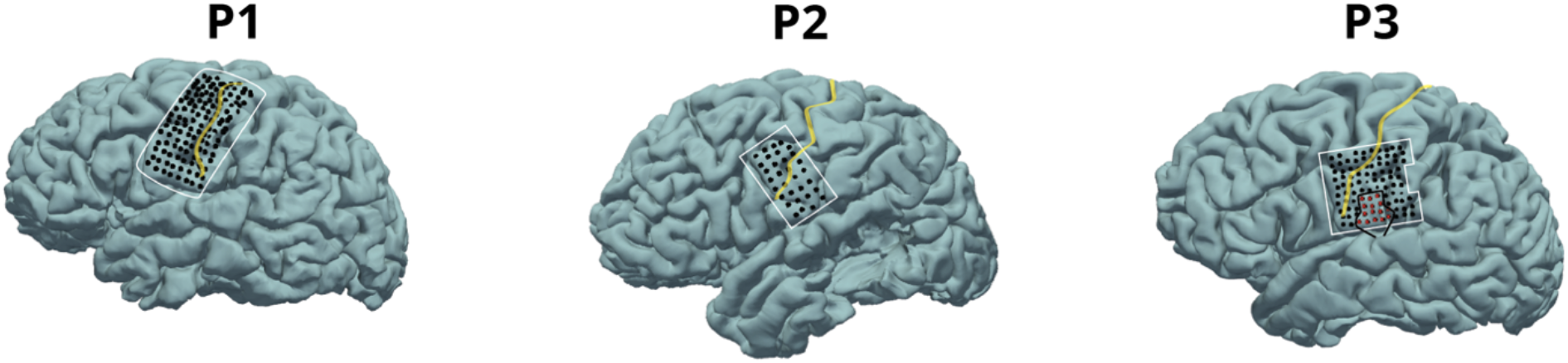
Locations of high-density (HD)-ECoG grid electrodes. High-density ECoG grids were implanted over the sensorimotor cortex on the left hemisphere. In P3, the HD grid was partly on top of another small grid blocking signals. These electrodes were removed from all analyses (marked by red crosses). The yellow lines indicate the location of the central sulcus.

HD-ECoG signals were recorded at a sampling rate of 2000 Hz for P1 and P3 (Blackrock Microsystems LLC, Salt Lake City, USA), and at 2048 Hz for P2 (Micromed, Treviso, Italy). For P1 and P3, audio recordings were recorded by a microphone connected to the neural recording system during the task. For P2, audio recordings were obtained with a separate microphone installed in the participant’s room. All audio recordings were recorded at a sampling rate or 30000 Hz. For P1 and P3, the audio was recorded by Blackrock and was automatically synchronised with brain data. For P2, the audio was recorded by the Presentation^®^ software (Version 23.0, Neurobehavioral Systems, Inc., Berkeley, CA, www.neurobs.com) on the stimulus presentation computer and was manually synchronised with brain data based on the Presentation log.

#### Experimental procedures

Presentation^®^ software (Version 23.0, Neurobehavioral Systems, Inc., Berkeley, CA, www.neurobs.com) was used to run the experiments for P1 and P2, and PyQT (https://wiki.python.org/moin/PyQt) was used to run the experiment for P3. As in the EMA experiment, all HD-ECoG participants were instructed to read out the same 97 Dutch words, including 70 object names from the picture naming task and 27 CVC sequences from the CVC sequence reading task. However, unlike the picture naming task in the EMA experiment, all words in the HD-ECoG experiment were presented to participants in textual form to avoid visual responses evoked by pictures (Figure 1b). All words were presented in random order and each word appeared on the screen twice. The duration of word trials and the inter-trial interval varied across participants as P3 performed the task in the operating room, where time for research was limited. Thus, in P1 and P2, the duration of a word trial was 2 seconds, and the duration of an inter-trial interval was 1.2 seconds. In P3, both the word trial and the inter-trial interval had a duration of 1.8 seconds.

### Data Analysis

The data analysis framework (Figure 4) consists of two pipelines: the EMA analysis pipeline and the HD-ECoG analysis pipeline. Each pipeline includes three steps: data preprocessing, tensor construction, and feature extraction. From these steps, we obtained articulatory features extracted from the EMA tensor and neural features extracted from the HD-ECoG tensor. We then used these features for the articulatory feature reconstruction.

**Figure 4.**
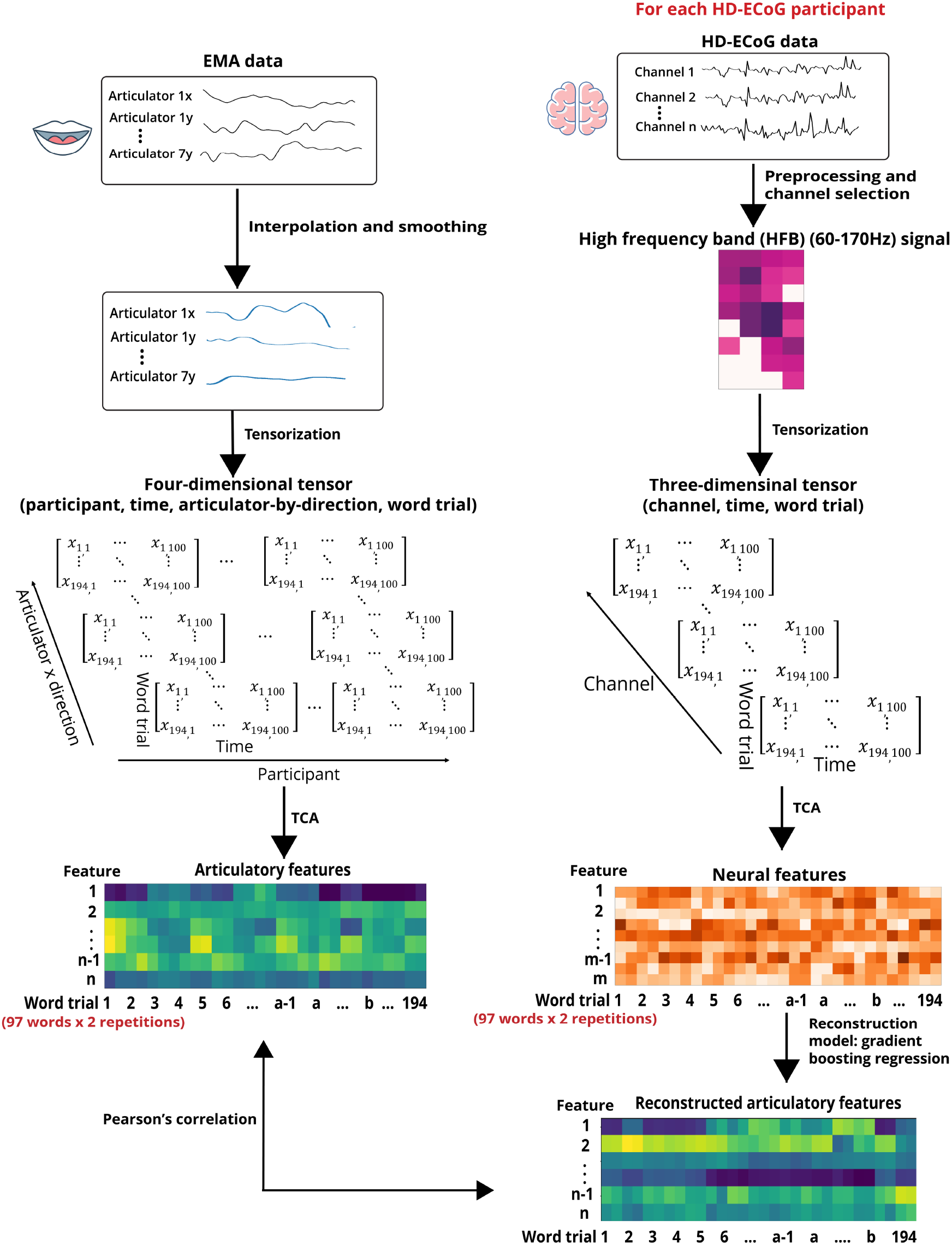
The framework for articulatory feature reconstruction from HD-ECoG signals. The left panel shows the analysis pipeline of the EMA and the right panel shows the analysis pipeline of the HD-ECoG dataset.

#### EMA data preprocessing and tensor construction

First, we identified missing values in the EMA data for each participant (U1: 1.35%, U2: 1.14%, U3: 0.16%, U4: 0.41%, U5: 0.71%, U6: 0.35%, U7: 0.42%, U8: 0.51%). Missing values were imputed using linear interpolation available in the Pandas library of Python (https://pandas.pydata.org). Then, EMA data were smoothed with a 5th-order, 20 Hz low-pass Butterworth filter. After that, EMA data in the left-right direction (z-axis) were excluded. We then removed baseline from the remaining EMA data by subtracting the mean positions for each combination of articulator and direction during a separate 10-second rest period, in which participants were asked to refrain from speaking or swallowing (Wieling *et al*., 2016).

After preprocessing, we segmented EMA data by using the time window from 0 to 1 second around the speech onset for each word trial and obtained a three-dimensional tensor: 100 x 14 x 194, which corresponds to 100 time points, 14 combinations of articulators (7) and directions (2), and 194 word trials (97 unique words repeated twice) for each participant. We then concatenated these tensors across participants and obtained a four-dimensional tensor (8 x 100 x 14 x194). We experimented with shifting the time window around the speech onset (prior to speech onset) and different window sizes, but the current time window contained the most information.

#### HD-ECoG data preprocessing, channel selection and tensor construction

The HD-ECoG data analysis pipeline was applied to each HD-ECoG participant’s dataset. First, we visually inspected if there were channels with noisy or flat signals in the data. No channel was removed during this step for any of the participants. We then removed 14 channels in P3 because of they lay on top of another grid and therefore were not recording signals directly from the brain tissue (Figure 3). Next, for each remaining channel, line noise (50Hz and its harmonics) was removed with a notch filter. Common average referencing was then applied to remove common noise and trends, and HD-ECoG signals were downsampled to 512 Hz. After that, for each channel high frequency band (HFB; 60-170 Hz) signals were extracted in 1 Hz bins using Morlet wavelet decomposition and were averaged across frequency bins. HFB signals were then log-transformed and downsampled to 100 Hz to match the sampling rate of EMA. Finally, for each channel, the resulting HFB signals were baseline-corrected by subtracting the mean values of a 10-second rest period before the first word trial.

The speech onset moments were detected by Azure speech-to-text service (https://learn.microsoft.com/en-us/azure/ai-services/speech-service/fast-transcription-create?tabs=locale-specified) and manually corrected using Praat (Boersma, 2001). We used the time window from 0.25 sec before to 0.75 sec after speech onset to segment continuous recordings into word trials. The window starting at 0.25 s before speech onset captured all information of the first phoneme in the sensorimotor cortex and has been shown to produce optimal decoding results in previous studies (Jiang *et al*., 2016; Ramsey *et al*., 2018). The 1 second window length captured the articulatory movements of most words (95 percentile of the distribution of word durations: 0.69 sec). Rest trials were composed of 1 second of data starting after the end of the preceding word trial. We identified channels with significant responses to the task by comparing the mean HFB activity between speech and silence using a two-tailed paired t-test. Since most channels showed very significant p-values (p<0.001), we selected channels with a large effect size (Cohen’s d >0.8, corresponding to a t-value of 12.44) to ensure only the most robust task-specific activation were included. The selected channels are referred to as active channels in subsequent analysis.

After channel selection, we constructed a three-dimensional tensor (100 x the number of selected channels x 194) for each HD-ECoG participant. Unlike the EMA analysis, we did not concatenate the HD-ECoG tensors across participants to construct a four-dimensional tensor, as the channel dimension differed across participants in number, location and electrode spacing (Figure 3 and Table 1).

#### Feature extraction

##### Tensor component analysis

Tensor decomposition was applied to extract a set of matrices. Vectors within these matrices are referred to as components. Individual component captures the variability along each dimension. We analyzed the characteristics of these components quantitatively and qualitatively. The tensor decomposition and characteristics analysis are referred to as tensor component analysis (TCA) (Williams *et al*., 2018).

CANDECOMP/PARAFAC decomposition is one of the most commonly used tensor decomposition methods (Kolda & Bader, 2009). An n-dimensional tensor 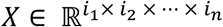 is decomposed into the combination of R components and can be approximated by,

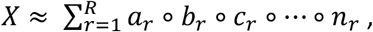

where ∘ is the outer product, and *a*_*r*_ ∈ *i*_1_, *b*_*r*_ ∈ *i*_2_, …, *n*_*r*_ (*r* = 1, …, *R*) are components, which are the r^th^ columns of the factor matrices *A, B*, … *N*, respectively. Elements in each factor matrix are factor loadings.

The CANDECOMP/PARAFAC decomposition procedure is often optimized by the alternating least squares method. This method has two steps. The first is to initialize factor matrices randomly or by certain criteria. The second step is to optimize one factor matrix at a time while keeping the others fixed. The optimization is achieved by minimizing the objective function between the original tensor and the approximated tensor. For example, when optimizing factor matrix *A*, we have the objective function,

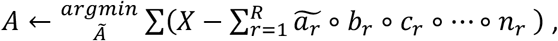

in which the minimization is a linear-square matrix problem. All factor matrices are optimized iteratively. When optimizing each factor matrix, this procedure is iterated until the objective function converges or the number of iterations reaches a pre-set threshold.

##### EMA tensor decomposition and articulatory feature extraction

We used the nonnegative CANDECOMP/PARAFAC decomposition optimized by hierarchical alternating least squares to decompose the EMA tensor. The decomposition pipeline was implemented using the Python library tensortools (Williams *et al*., 2018).

After applying the nonnegative CANDECOMP/PARAFAC decomposition, the EMA tensor was decomposed into 29 components and four factor matrices: the participant factor, time factor, articulator-by-direction factor and word trial factor. The decomposition process was repeated four times and resulted in four TCA runs with consistent results across the runs (Figure S3). We then checked the goodness-of-fit by inspecting the cumulative explained variance of all 29 components, which was 0.89 averaged across four runs (Figure 5a). Adding more components did not substantially increase the explained variance. To reduce components, we first selected components by the run with the highest explained variance. Since the explained variance of the first component was over 75%, we used the mean explained variance from the 2^nd^ to the 29^th^ component as the threshold to determine the number of components to select. The 10^th^ component was the last component with an explained variance above this threshold. Therefore, we only retained the first 10 components (Figure 5b). In the next step, our goal was to select components that generalized across all participants. To measure the generalizability of each component, we calculate the *commonality* value of each component in the participant factor as following (Hurley & Rickard, 2009; Williams *et al*., 2018),

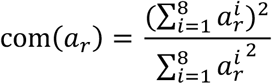

**Figure 5.**
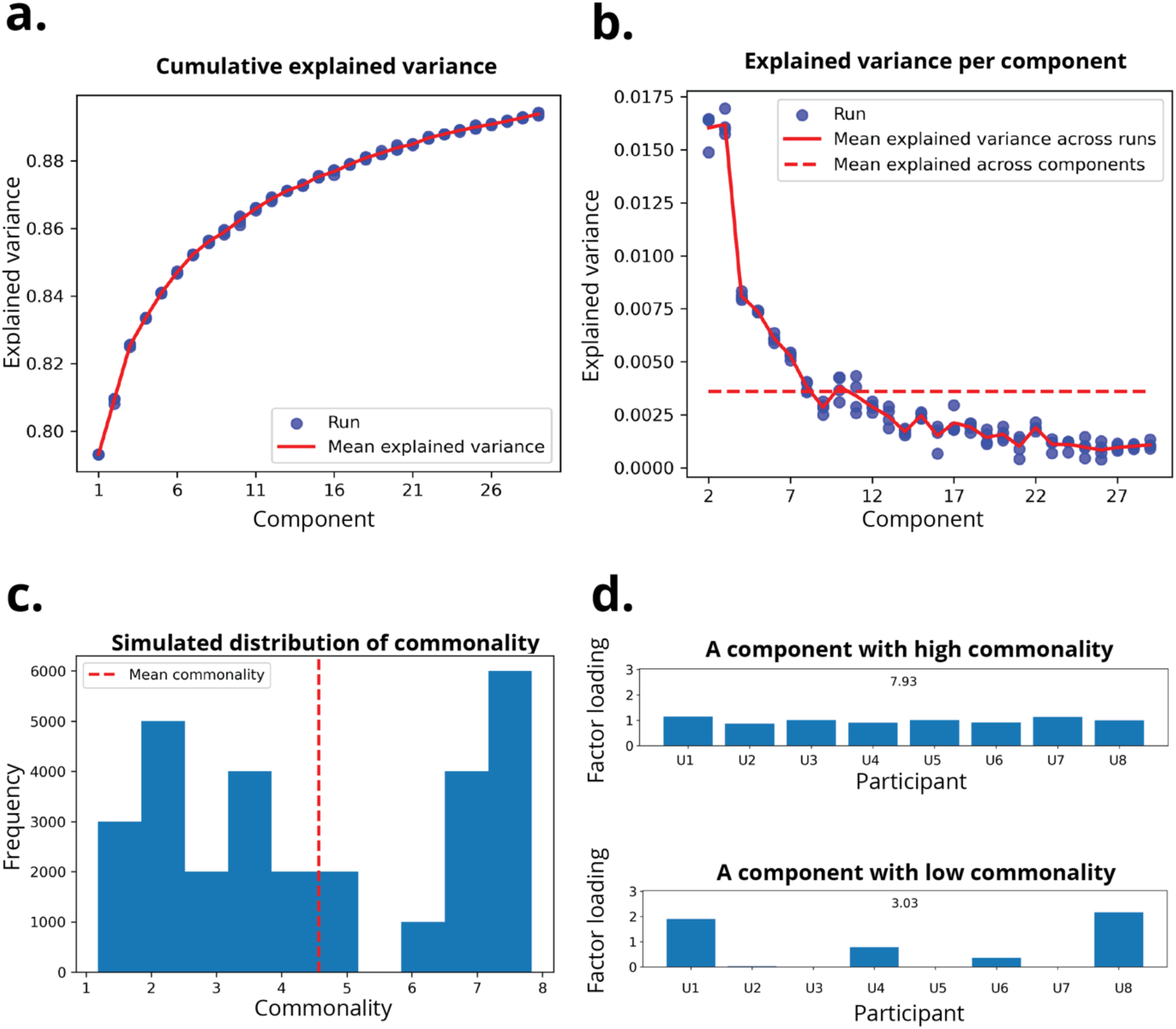
Tensor component selection in the EMA dataset. **a**. Cumulative explained variance of all 29 tensor components averaged across 4 runs. **b**. Explained variance per component (starting from component 2 since component 1 explained approximately 80% explained variance) averaged across 4 runs. The mean explained variance was indicated by the red line and used as the threshold for component pre-selection. Component 10th is the last component with an explained variance above the threshold. **c**. The simulated distribution of commonality obtained by permuting factor loadings across the 29 components. The red dotted line indicates the mean commonality of the simulated distribution. **d**. Examples of a component with high commonality and a component with low commonality, which was 7.93 and 3.03, respectively

In which *a*_*r*_ is the r^th^ component in the participant factor matrix and 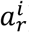 is the i^th^ element in that component. The commonality value reflects how many participants have similar factor loadings, i.e., how many participants contribute similarly to a component. If all participants have similar factor loadings, the commonality of this component equals the number of participants. In contrast, if only one participant has a non-zero factor loading, the commonality of this component equals 1, indicating that only that participant contributes to this component. We permuted elements across components 1000 times to simulate the distribution of commonality (Figure 5c). Figure 5d shows examples of a component with high commonality and a component with low commonality. We used the mean commonality (4.57) as the threshold, as this value indicates that more than half of eight participants contributed equally to the component. Using a higher threshold yields too few features and could compromise reconstruction performance. By using this threshold, six components with commonality above it were selected for further analyses.

We refer to selected components in these factor matrices as features. Specifically, in our reconstruction analysis, we refer to features from the word trial factor as the articulatory features.

##### HD-ECoG tensor decomposition and neural feature extraction

As in the articulatory feature extraction, we extracted features from the HD-ECoG tensors by using the nonnegative CANDECOMP/PARAFAC decomposition optimized by hierarchical alternating least squares. The HD-ECoG tensors were decomposed into 10, 14, and 17 components for P1, P2, and P3, respectively, corresponding to 80% explained variance. We chose 80% explained variance as the cut-off point for model fitting because adding more components did not increase the explained variance substantially.

To select reproducible features across repetitions, we used Pearson’s correlation coefficient (PCC) between word trial factor loadings to select components. Since HD-ECoG tensor decomposition yield less components than EMA tensor, only one-step component selection is needed. First, we shuffled word trial loadings across components 1000 times and calculated PCC between repetitions for all components. Figure 6 shows the distribution of PCC of word trial factors between repetitions for each HD-ECoG participant. We calculated the 95^th^ percentile of the distribution and used it to select components with PCC above this threshold from the word trial factor as neural features. This resulted in 7, 6, and 6 components for participants P1, P2, and P3, respectively. The mean PCC of extracted components between repetitions was 0.59 for P1, 0.30 for P2 and 0.49 for P3.

**Figure 6.**
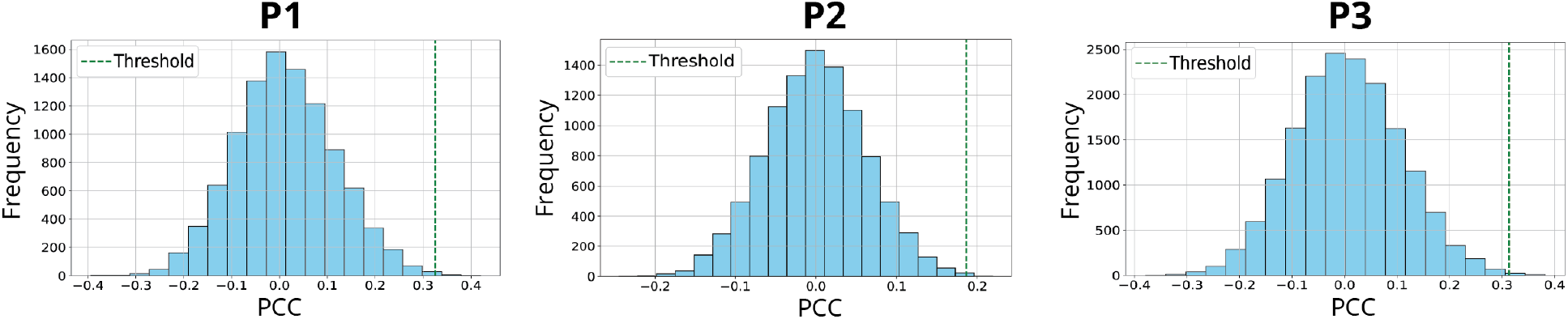
Selection of neural features in the HD-ECoG data. The panels show the distributions of PCC of word factors between word repetitions across all components over 1000 shuffles for HD-ECoG participants P1, P2, and P3, respectively. The significance threshold (α=0.05) (indicated by the green dotted line, which represents the 95^th^ percentile of the distribution) of each distribution was 0.33, 0.17, and 0.31 for participant P1, P2 and P3, respectively. We selected components with a PCC above these thresholds.

As in the EMA analysis, we combined vectors from the selected components to create a factor matrix for each dimension of the HD-ECoG tensor. We obtained three factor matrices: the channel factor, the time factor, and the word factor, which captured spatial, temporal, and kinematic variations across words in neural activity, respectively. We also refer to components from these factor matrices as features. In our reconstruction analysis, features from the word trial factor matrix are referred to as the neural features.

### Generalisable articulatory feature reconstruction from neural features

To capture the complex relationships between neural data and articulatory data, we used the gradient boosting regression model to reconstruct the generalisable articulatory features from the neural features (Friedman, 2001; 2002; Hastie, 2009). This model was chosen for its robust performance on small datasets so that overfitting can be avoided. Reconstruction performance was evaluated by the PCC between the reconstructed and original articulatory features and was cross-validated across all words using the leave-one-word-out scheme. We used the permutation test to determine the significance of reported PCC values by shuffling word labels of all trials 1000 times. Permutation was performed on data of repetition 1 and repetition 2 separately.

## Results

### HD-ECoG channel selection

For P1, P2, and P3, respectively, 19.5% (25/128), 75% (24/32) and 25.6% (21/82) of HD-ECoG channels showed very strong HFB increases during word trials. These channels were referred to as active channels (Figure 7).

**Figure 7.**
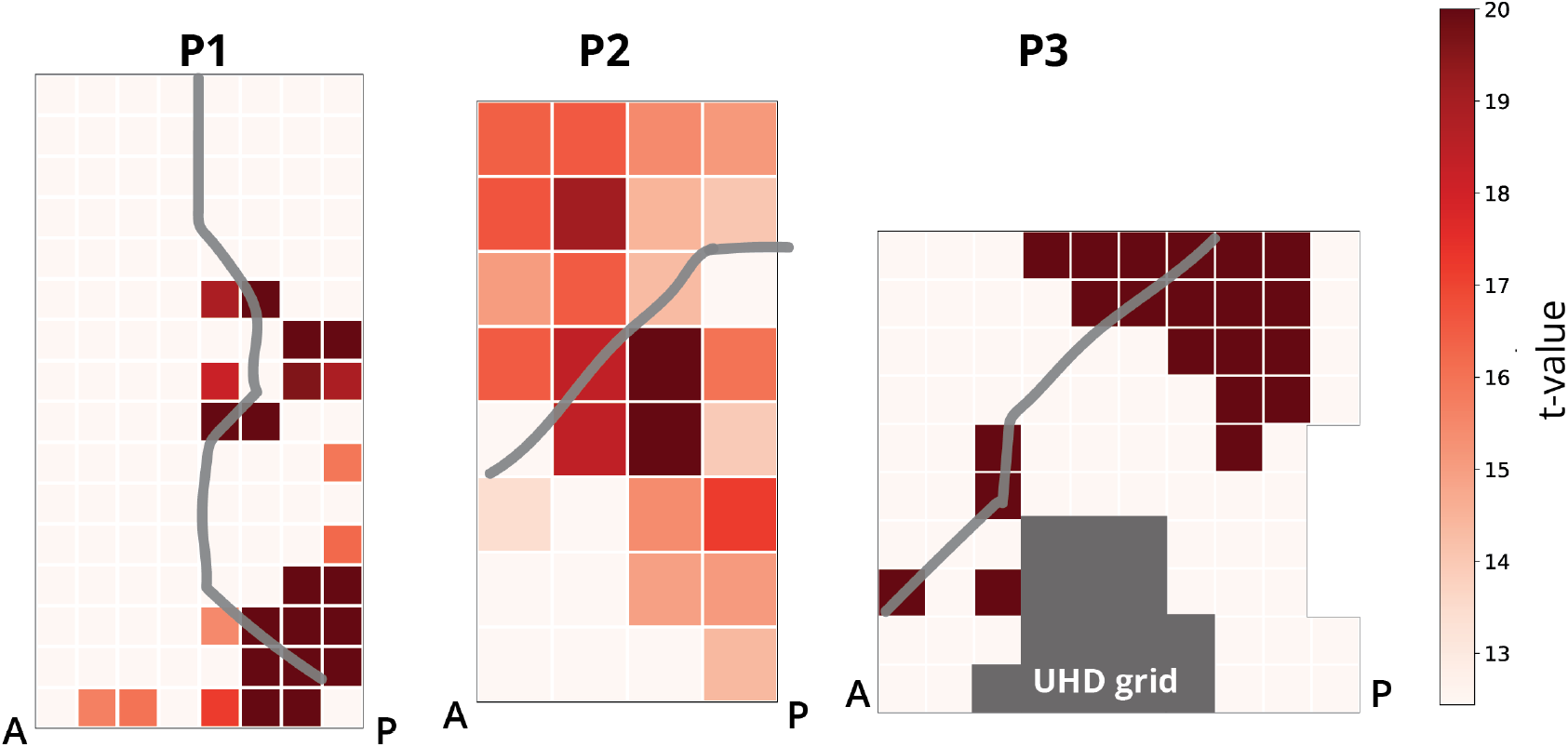
T-value maps comparing high-frequency band (HFB) responses during speech periods and silence periods. The channel colour represents t-values, as shown in the colormap. The threshold for t-values is 12.44, corresponding to a large effect size (Cohen’s d>0.8). Only channels with a t-value above the threshold were color-coded to highlight the most robust task-specific activation. In P3, the greyed-out channels were lay on top of another grid. The grey line indicates the location of the central sulcus. Label “A” represents the anterior direction and label “P” represents the posterior direction.

### Features extracted from the EMA tensor

Figure 8 shows the time, articulator-by-direction, participant and word trial factors extracted from the EMA tensor. The time factor shows temporal profiles of kinematic patterns shared across all articulators. The articulator-by-direction factor illustrates the strength of movements per articulator along different directions. Considering both factors together reveals how the kinematic patterns evolve across time for each articulator. For example, features 2 and 3 capture the tongue movements along the anterior-posterior direction within the first 0.5 seconds after the speech onset. Overall, most movements occurred within 0.5 sec after speech onset, which is the mean duration of word production.

**Figure 8.**
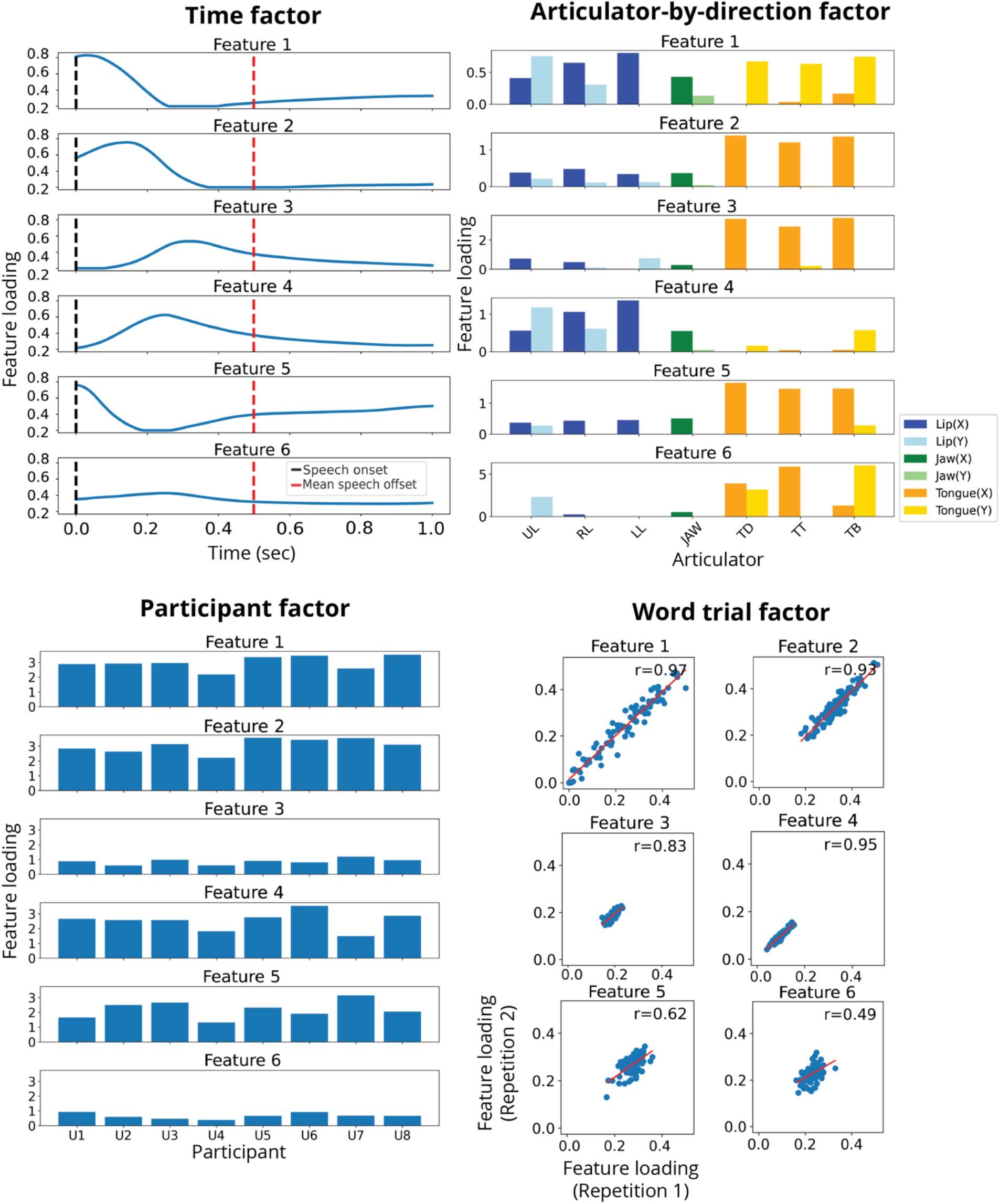
Articulatory features extracted from EMA data using TCA. T**he participant factor of the EMA tensor**. Each bar represents the contribution of each participant to the feature. **The articulator-by-direction factor of the EMA tensor**. Each bar shows movements recorded by a sensor along anterior-posterior (x) or up-down (y) direction. We grouped sensors into three groups by articulators (lip, jaw, and tongue) and colored them in blue, green and orange according to the group. The dark colors (dark blue, dark green and dark orange) represent movements along the anterior-posterior (x) direction, and the light colors (light blue, light green, and light orange) represent movements along the up-down (y) direction. **The time factor of EMA tensor**. The black line indicates the speech onset, and the red line indicates the mean duration of spoken words averaged across all trials and across all participants. **The word factor of the EMA tensor**. We showed the Pearson’s correlation (r) between factor loadings of repetitions for all words, as well as the linear regression fitted line between repetitions (shown as the red line).

In the participant factor, the commonality values of features 1-6 were 7.92, 7.88, 7.39, 7.32, 6.37, and 4.72, respectively, indicating that over half of participants contribute similarly to the selected features. This result means that the extracted features generalize well across participants. The word trial factor reflects the spatiotemporal articulatory features for each word trial. In the word trial factor, the PCC between repetition 1 and 2 were 0.97, 0.93, 0.83, 0.95, 0.62, and 0.49. PCC values were significant (P<0.05) for the first four features. Features 5 and 6 show a lower PCC, as well as lower commonality. Taken together, features 5 and 6 may reflect trial-specific articulatory kinematics patterns.

### Features extracted from HD-ECoG tensor

As in the EMA tensor, each dimension in the HD-ECoG tensor corresponds to one factor. Supplementary Figure S1 show the time, channel factor and word factor for P1, P2, and P3. Features in the time factor show the time profiles of basis functions that reflect the HFB activity shared across all active channels. In all participants, we identified three types of brain responses by inspecting the time profiles: early response, mid response and late response. For each participant, the basis function reaching its peak amplitude prior to speech onset is marked as the early response; the basis function showing rising activity after speech onset and reaching its peak amplitude before mean speech offset is marked as the mid response; and the basis function reaching its peak amplitude around or after the mean speech offset is marked as the late response (Figure 9). Features in the channel factor illustrate how the basis functions in the time factor are weighted for each channel, suggesting the anatomical localization of basis functions. In P1 and P3, channels with large weights are most located posteriorly, while in P2, channels with large weights are most located anteriorly. When examining the anatomical localization of different types of brain responses, we did not observe a clear distinction in P1 and P3. However, in P2, the early response is evenly located across all active channels, while the mid and late responses are more anteriorly located.

**Figure 9.**
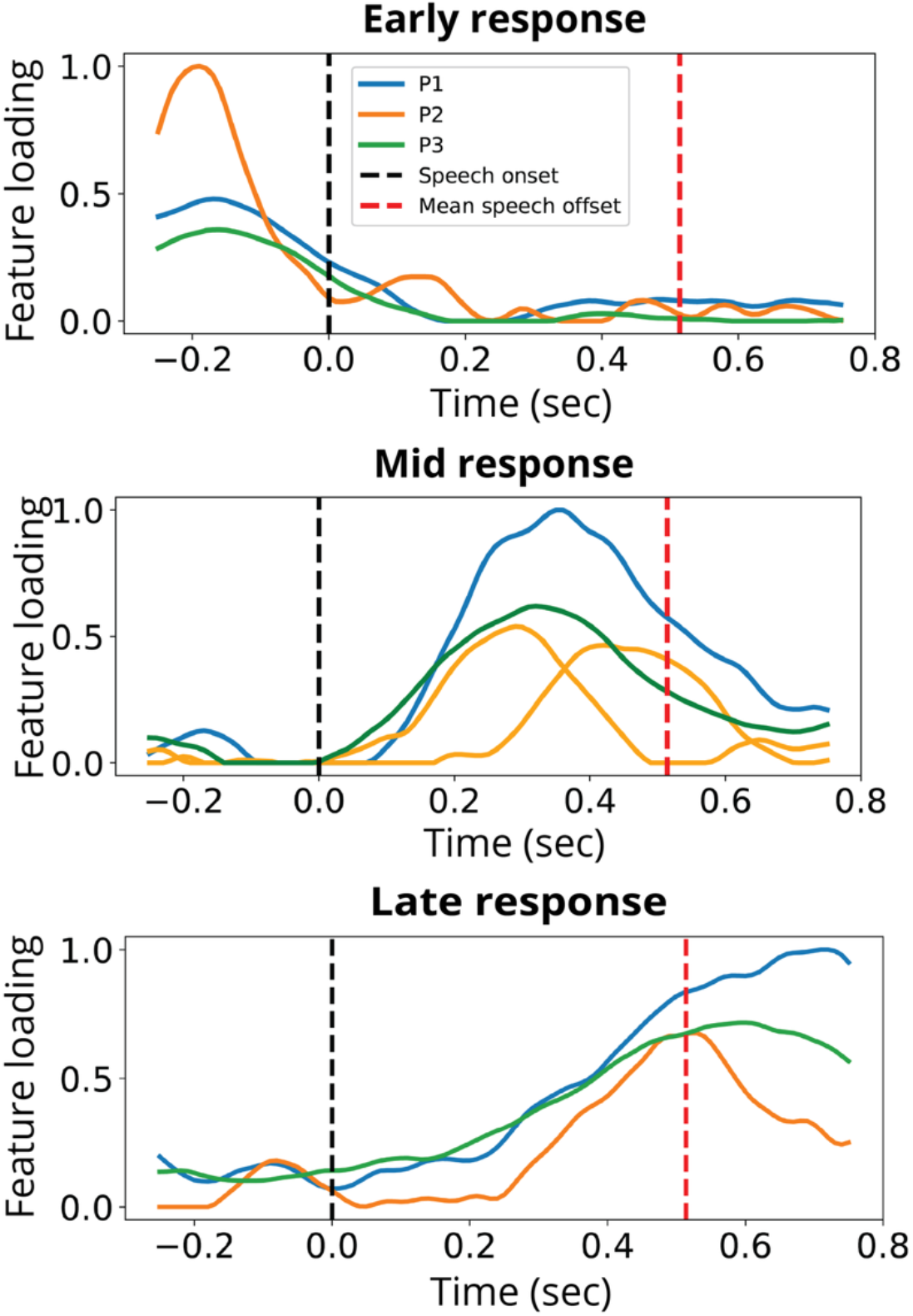
Three types of brain responses during the task: early, mid response and late responses. All temporal profiles were min-max normalized to enable direct comparison across participants. The early, mid, late response are shown by the blue, orange and green line, respectively. We identified the complete set of time features for each response type. For the early and late response, only one time feature was identified per participant, whereas for the middle response, two time features were identified in P2. The speech onset is shown by the black dashed line, and the mean speech offset by the red dashed line.

Features in the word trial factor reflect the spatiotemporal neural features for each word trial, i.e., the combination of anatomical distribution of neural activation and time profiles of brain response. The PCC corresponding to early responses appeared to be lower than those corresponding with mid and late responses.

### Generalisable articulatory feature reconstruction

We reconstructed articulatory features from neural features by using the gradient boosting regression model. Figure 10 demonstrates the reconstruction performance averaged across word repetitions. Among three HD-ECoG participants, P1 achieved the best reconstruction performance, with a mean PCC of 0.80 (p<0.05), while P2 and P3 achieved a mean PCC of 0.75 (p<0.05) and 0.76 (p<0.05), respectively.

**Figure 10.**
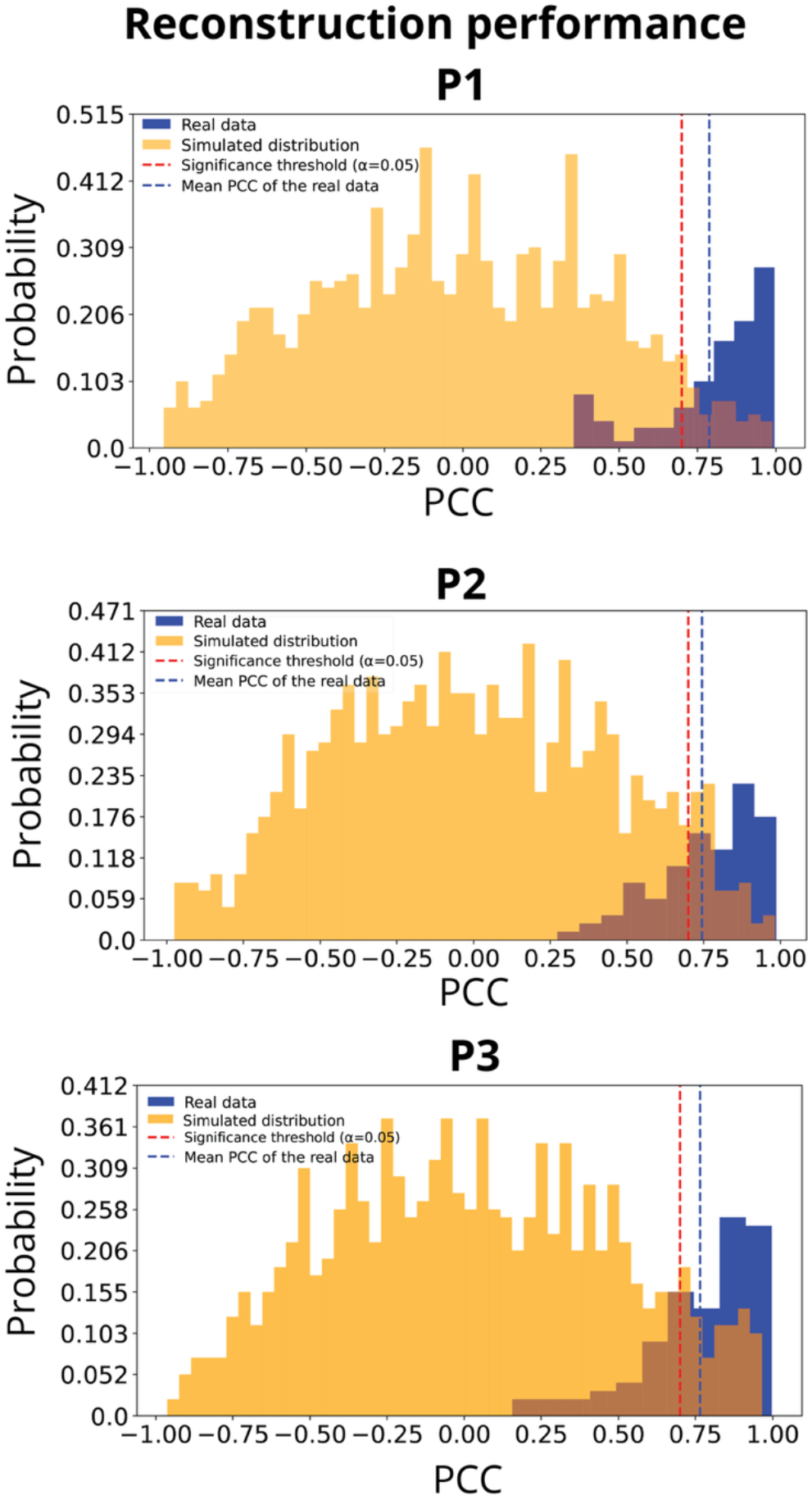
Results of articulatory feature reconstruction from neural features. For each participant, we calculated the real distribution of PCC (shown in blue). We also calculated the simulated distribution of PCC by shuffling word labels of all features 1000 times (shown in yellow). The red dashed line denotes the significance threshold (α = 0.05), and the blue dashed line denotes the mean PCC of the real distribution.

## Discussion

In our study, by recording brain activity from the vSMC using HD-ECoG grids and leveraging a publicly available EMA dataset, we demonstrated a generalizable BCI framework that reconstructed articulatory features from three participants without relying on their articulatory or acoustic data. For individuals with severe vocal tract paralysis, this generalizable BCI framework, which is independent of individual-specific articulatory or acoustic data, could be a promising way to restore speech from brain activity.

Because articulator movements were not measured in the HD-ECoG participants, we used a TCA model to extract articulatory features from a public EMA dataset. We then selected features with high commonality values, indicating that these articulatory features capture articulatory movement patterns shared across speakers. Earlier studies also attempted to align articulatory features across speakers. For example, Bocquelet and colleagues used a linear model to map the articulatory features of individual speakers to the feature space of a reference speaker (Bocquelet *et al*., 2016). However, their approach can be biased by the articulatory characteristics of the chosen reference, such as vocal tract shape. In contrast, our approach employed commonality to ensure that the selected features showed generalizable articulatory kinematics across healthy speakers, which is particularly advantageous when both acoustic and articulatory data cannot be obtained from target users. A previous study not having articulatory movement data available reported good results using an acoustic-to-articulatory inversion model relying on participants’ own audio data to generate articulatory data for speech BCI development (Anumanchipalli *et al*., 2019). However, their approach requires participants to produce intelligible speech, which is not feasible for individuals with severe vocal tract paralysis.

Having extracted a set of generalizable articulatory features across speakers, we next examined the interpretability of the extracted neural features. In all three participant we identified three types of brain responses: early, mid and late. Compared with previous studies, the early and late responses we observed resemble the transient response, which shows increased activity around speech onset/offset, and the mid response resembles the sustained response, which shows increased activity during the utterance (Conant *et al*., 2018; Salari *et al*., 2018). However, we did not observe a clear distinction in the anatomical organization among these responses, while Salari and colleagues found that transient responses were more anteriorly located and the sustained responses were more posteriorly located in their participants with HD grids (Salari *et al*., 2018). One possible explanation is the relatively small number of active channels in our analysis. Beyond the anatomical organization of these brain responses, we also considered their possible functional roles: early and late responses may reflect the articulator movements at the speech onset/offset, such as mouth opening or closing. Alternatively, the early response may also reflect movement planning (Salari *et al*., 2018). For the mid response, it may represent a nonspecific signal for holding the vocal tract configuration (Conant *et al*., 2018). To further understand the neural representation during word production, more HD-ECoG data will be needed.

To reconstruct articulatory features from brain activity, we trained a gradient boosting regression model using word trial factors from both the EMA and HD-ECoG TCA models. In all three participants, the reconstructed and the original articulatory features significantly correlated with each other. However, the correlation coefficients were only slightly above the significance threshold (α = 0.05), indicating that the articulatory features may be similar across words. One possibility is that the articulatory features of different words cluster closely in the low-dimensional space. Further studies should include more repetitions per word and train a word-level classifier to evaluate the discriminability of articulatory features.

To the best of our knowledge, this is the first study to extract generalizable articulatory features from healthy individuals and reconstruct them from brain activity of different individuals. Previous work showed that speakers share a similar state-space of articulatory features, but only used this to reconstruct voice, demonstrating the cross-subject articulatory-acoustic transfer (Anumanchipalli *et al*., 2019). Our study extends their work by showing the cross-subject articulatory-neural transfer, demonstrating that the generalizable articulatory features can be reconstructed from brain activity, even if the two modalities are from different individuals. This finding is also consistent with prior research, where authors report structural similarity between the articulatory features from healthy participants and neural features from an individual with paralysis (Willett *et al*., 2023). Such cross-subject articulatory-neural transfer is beneficial for speech BCI design: even for individuals with severe vocal tract paralysis, it may still be possible to reconstruct articulatory movements by developing models using articulatory data from healthy individuals.

Recent advances in speech BCIs have shown the possibility of reconstructing speech directly from brain signal without relying on intermediate articulatory features (Wairagkar *et al*., 2024; Littlejohn *et al*., 2025). However, previous research suggests that articulatory features are more robustly encoded in the vSMC than acoustics features, and therefore can be learned faster with limited neural data (Chartier *et al*., 2018; Conant *et al*., 2018; Anumanchipalli *et al*., 2019). Thus, incorporating intermediate articulatory features may enhance the speech decoding performance (Anumanchipalli *et al*., 2019). Apart from audio synthesis, the intermediate articulatory features can also be used to restore speech-related orofacial movements (Metzger *et al*., 2023).

Taken together, our results indicate that our generalizable BCI framework could be a promising approach for restoring the communication for individuals with severe vocal tract paralysis, for whom articulatory or acoustic data are not available. However, our current use of EMA data, which only contains 7 sensors attached to the upper vocal tract, restricts the ability to capture complete vocal tract movements. Using more complete articulatory features that span the entire vocal tract space may enable speech synthesis with human-like fidelity (Wu *et al*., 2023).

Another limitation of our study is the vocabulary size. Our restricted vocabulary of 97 Dutch words is likely not enough to capture the full range of articulatory gestures in natural conversation. Expanding to larger vocabularies and to sentences will be important for assessing reconstruction performance in more realistic contexts. Finally, the proposed pipeline has only been validated offline. In the future, adapting this framework for online use could enable real-time articulatory-based speech BCIs.

## Conclusion

Overall, we demonstrated that our proposed framework could reconstruct generalizable articulatory features from brain activity from a separate group of able-bodied speakers, even when these speakers’ articulatory or audio data were not available. Using these generalizable articulatory features has potential for developing speech BCIs that can restore full communication for individuals with severe vocal tract paralysis, while reducing the need for a large amount of training data.

## Acknowledgement

This work is supported by Dutch Brain Interface Initiative (DBI2), project number 024.005.022 of the research programmed Gravitation, which is financed by the Dutch Ministry of Education, Culture, and Science (OCW) via the Dutch Research Council (NWO).

## Supplementary materials

**Figure S1.**
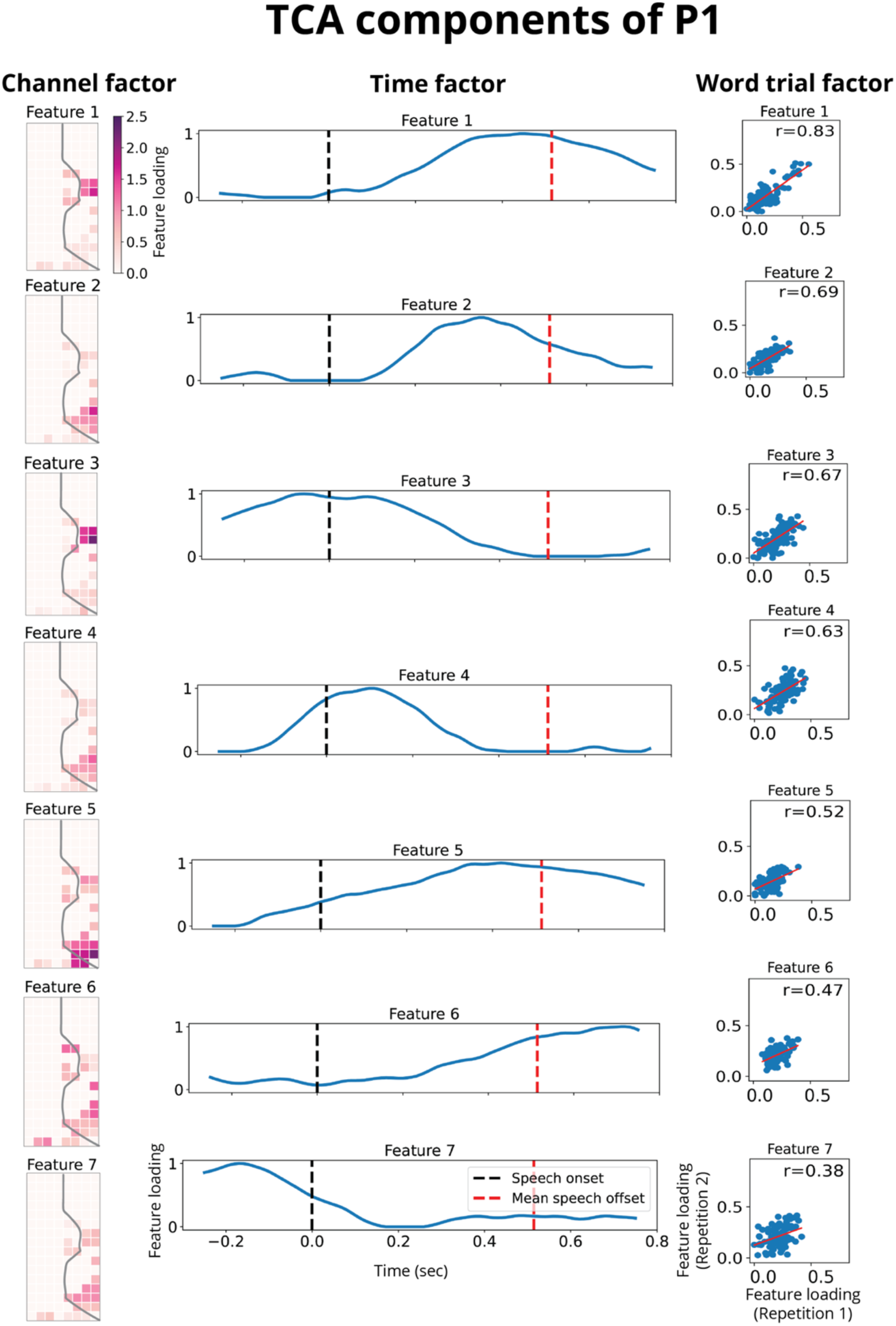
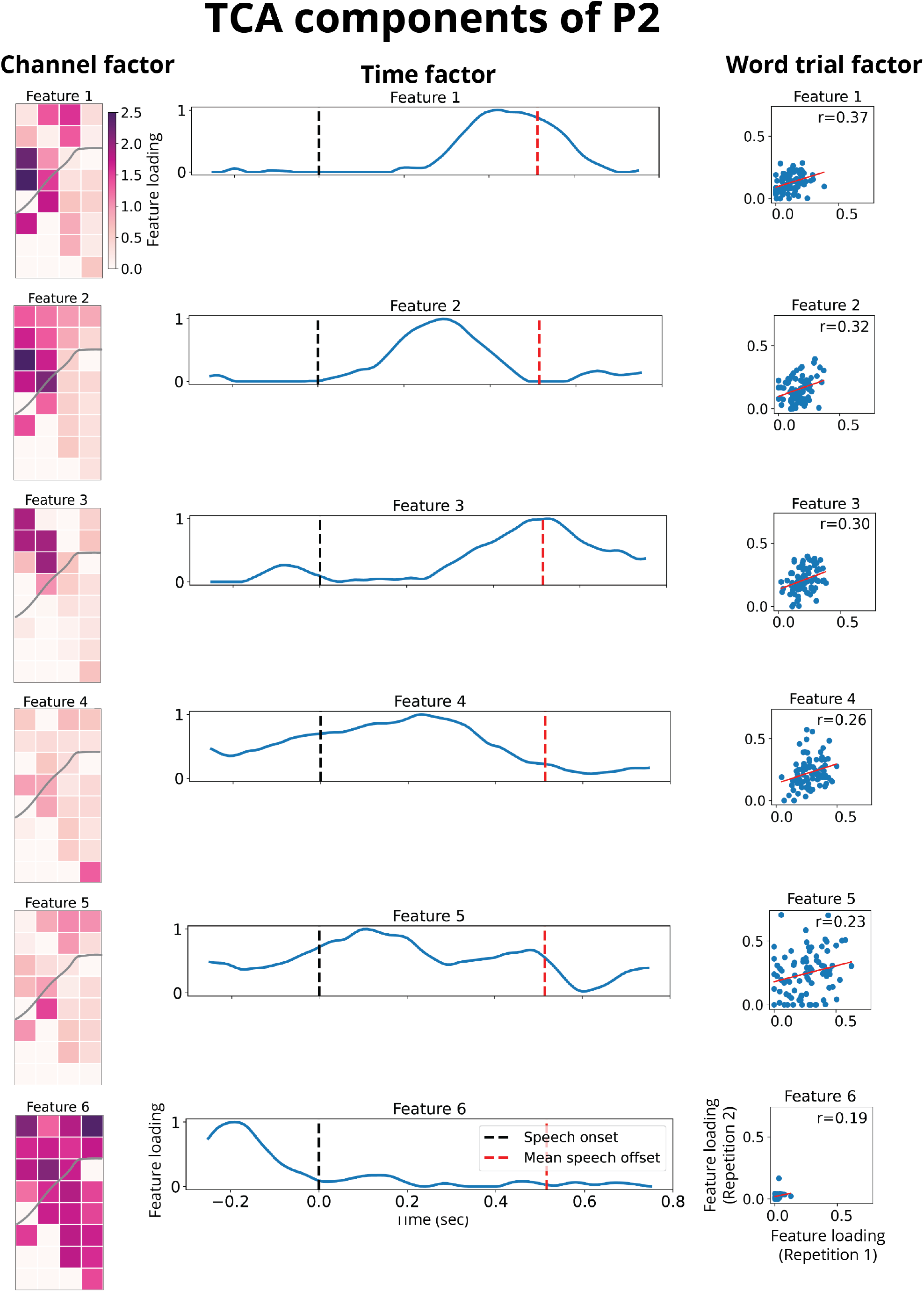
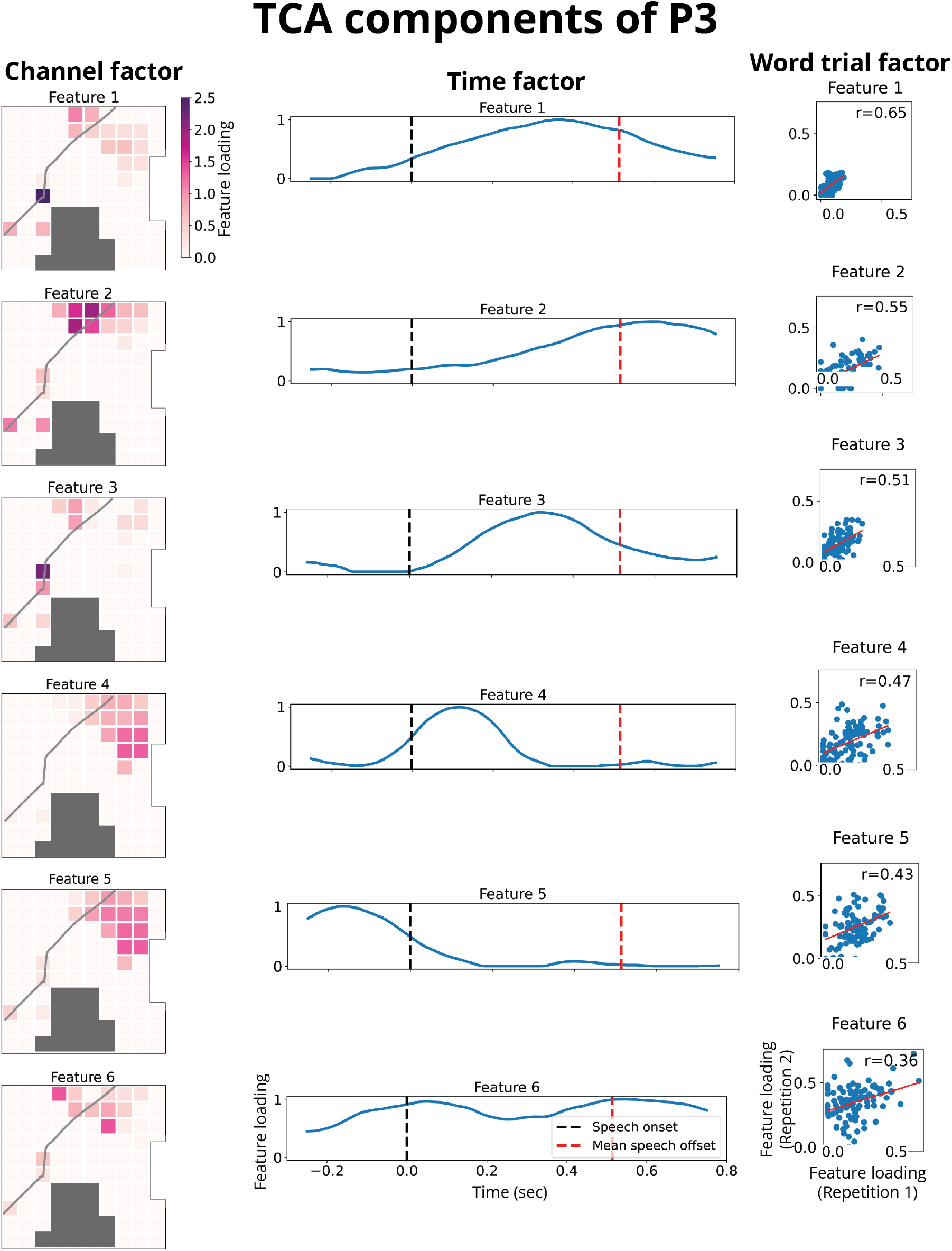
ECoG Components of P1, P2, and P3. From left to right in each panel, we showed the channel factor, the time factor, and the word factor for participant P1, P2 and P3, respectively. In the channel factor, the grey lines indicate the location of central sulcus. In the time factor, the black dash lines indicate the speech onset and the red dash lines indicate the speech offset averaged across word trials. For components with the flat line, no brain activity was reflected during the corresponding period. In the word factor, the red lines are the linear regression fitted line between feature loadings of repetition 1 and 2. We also showed the Pearson’s correlation (r) between feature loadings of repetition 1 and 2 for each selected feature.

**Figure S2.**
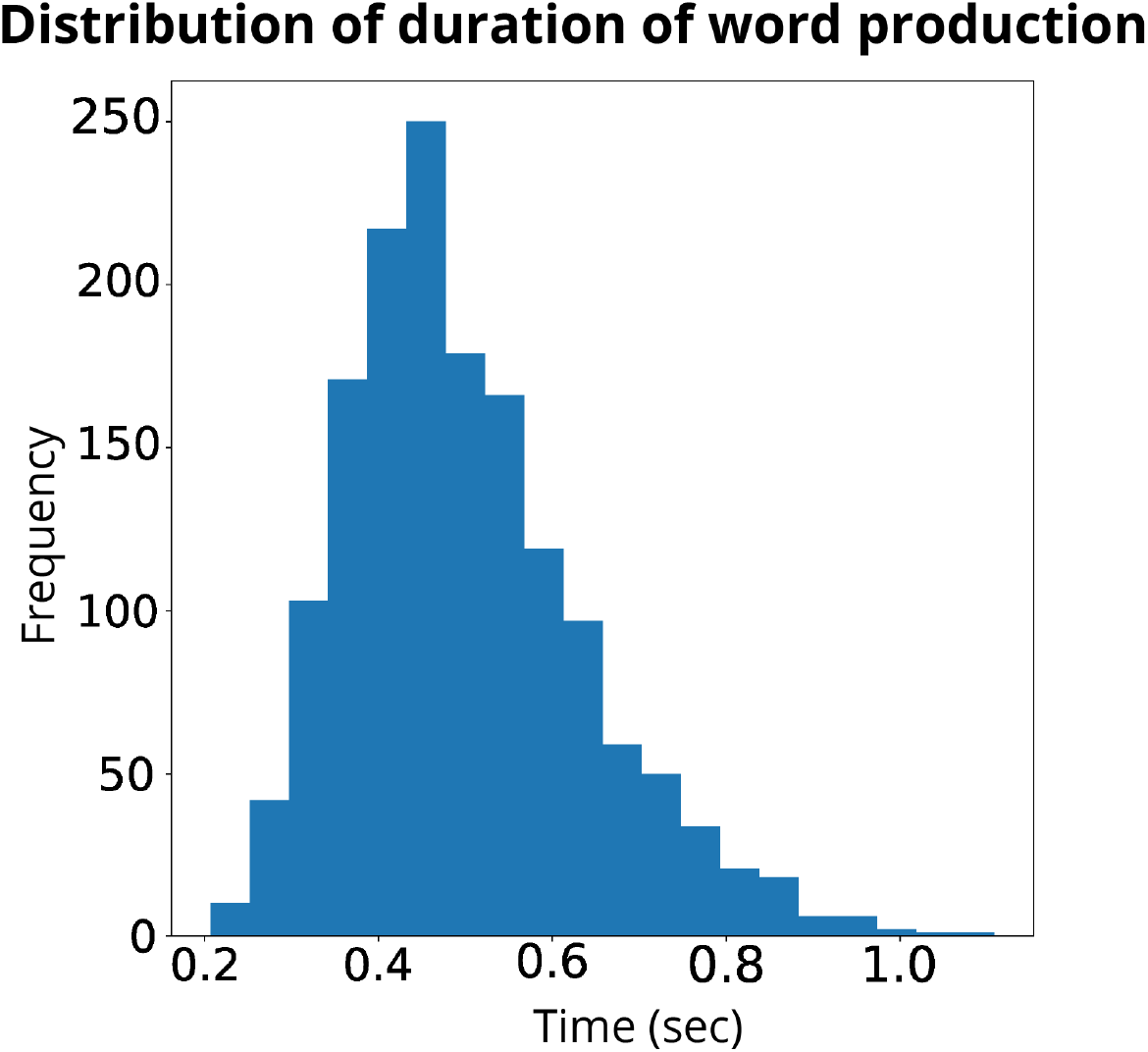
Distribution of word production during HD-ECoG data collection. We accumulate all word production duration, which is the interval between speech onset and offset across all word trials for three HD-ECoG participants and calculate the distribution.

**Figure S3.**
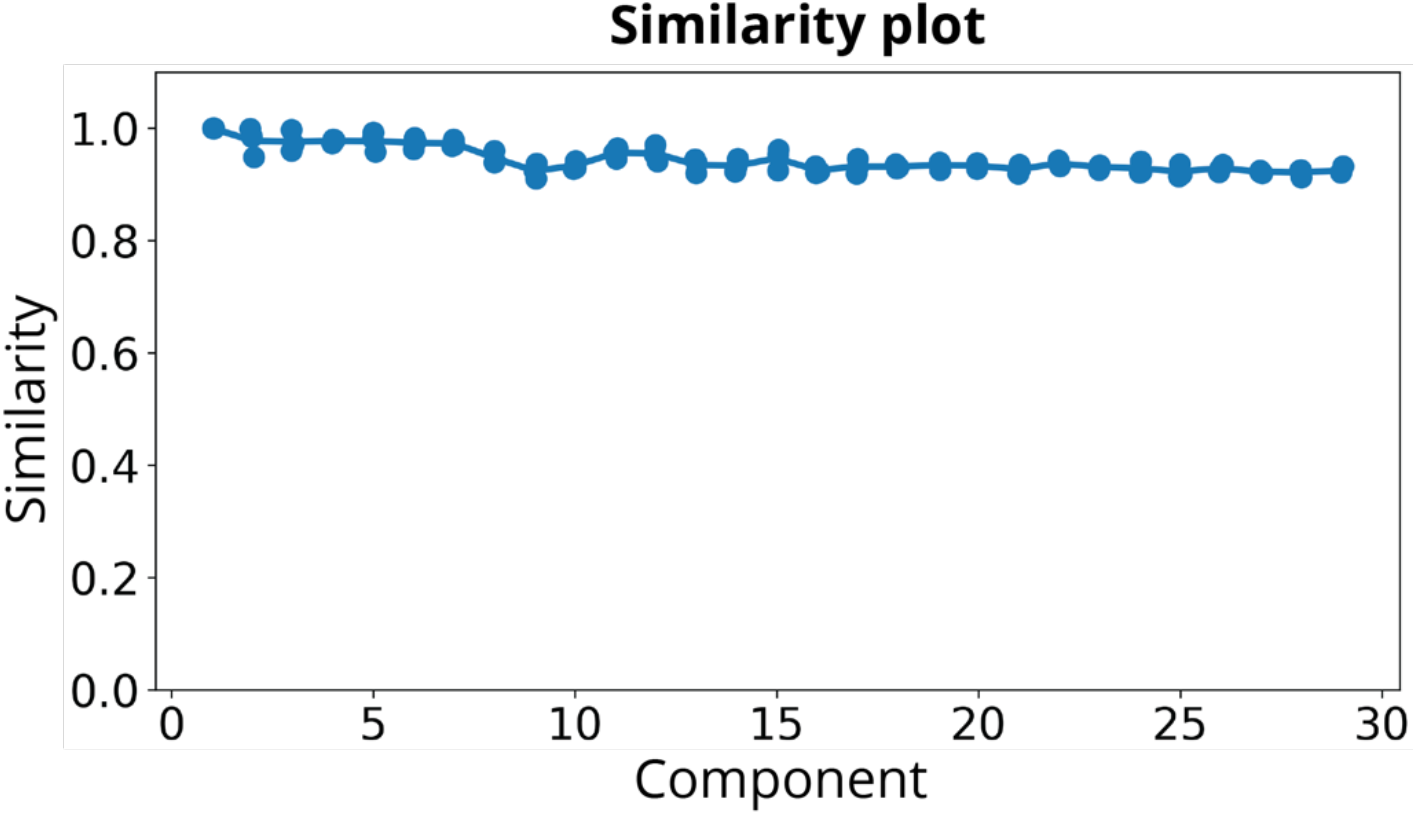
Similarity scores between TCA runs with different number of components. For each TCA run (blue dot), a similarity score was calculated across all TCA runs and the run with the lowest reconstruction error

**Table S1.** Average variance per articulator-by-direction across participants. For each articulator, the variance of movements along each direction: anterior-posterior (X), up-down (Y), and left-right (Z) was calculated and average across participants.

| Articulator \ Direction | Anterior-posterior (x) | Up-down (Y) | Left-right (Z) |
| --- | --- | --- | --- |
| UL | 1.41 | 1.62 | 1.52 |
| RL | 4.27 | 1.20 | 4.09 |
| LL | 6.18 | 4.38 | 3.19 |
| JAW | 2.23 | 2.55 | 1.69 |
| TD | 16.57 | 8.77 | 3.96 |
| TB | 17.22 | 9.46 | 3.93 |
| TT | 18.81 | 9.52 | 5.24 |

### Similarity score

The similarity of two fitted TCA models were calculated based on the angles between latent factors (Tomasi & Bro, 2006; Williams *et al*., 2018). For two fitted four-factor TCA models with R components, {*A, B, C, D*} and {*A*^*′*^, *B*^*′*^, *C*^*′*^, *D*^*′*^}, the similarity score is,

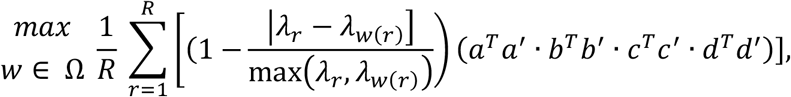

where Ω denotes all possible permutations of all factors, and *w* is a particular permutation. When calculating the similarity score, every time one component is selected from each of the four factors, each scaled to unit length and *λ*_*r*_ denotes the product of these scalings. The similarity score is averaged across all components. After enumerating all possible permutations, the maximal similarity score is taken as the final similarity score between these two TCA models.

